# Power and linkage disequilibrium differences are major confounders of human eQTL portability across ancestries and cohorts

**DOI:** 10.64898/2026.02.26.708346

**Authors:** Patrick M. Gibbs, Isobel J. Beasley, Christina Brady Del Azodi, Davis J. McCarthy, Irene Gallego Romero

## Abstract

The phenotypic effects of germline variants are often mediated through gene regulation. Expression quantitative trait loci (eQTLs) are genetic variants associated with changes in gene expression. Understanding how eQTLs vary across populations is essential for characterising the genetic and regulatory drivers of trait diversity. Meta-analysing eQTL studies from multiple populations enables more robust detection of eQTLs and can reveal regulatory mechanisms shaped by population-specific environmental or ancestry-related factors. However, across the multi-ancestry eQTL literature, a wide range of methods have been used to quantify eQTL portability across ancestry groups. Because different studies employ different portability metrics, it is challenging to form a coherent view of the regulatory landscape across populations. In this work, we analyse eQTL summary statistics from ten datasets matched on tissue type and sequencing technology. We compare portability metrics used previously and show that they can yield markedly different patterns of apparent regulatory conservation or divergence. We then examine the statistical determinants of portability across metrics and demonstrate that sample size, minor allele frequency, and linkage disequilibrium are major drivers of the observed differences in eQTL portability across studies. These findings highlight that differences in statistical power stemming from factors such as population size and allele frequency must be accounted for when evaluating eQTL portability. To address this issue, we introduce a new approach designed to correct for these factors when calling eQTL portability. Finally, we show that empirical Bayes multivariate adaptive shrinkage provides a powerful framework for meta-analysing multiple eQTL studies, with the ability to pool signals across populations to produce more robust effect-size estimates within each population, and better powered detection of population specific eQTLs.

**Author Summary:** Genetic variants often influence human traits and diseases by altering gene regulation. Expression quantitative trait loci (eQTLs) are variants associated with changes in gene expression. Many eQTLs are not portable — they fail to reproduce across cohorts and ancestries — yet studies use many different metrics to decide when an eQTL has reproduced, making it hard to form a coherent picture of gene regulation across populations. Here we compiled eQTL studies from ten cohorts spanning ancestries, tissues, and sequencing technologies, showing that commonly used portability metrics reveal distinct patterns of sharing. We find the main drivers of non-portability are differences in sample size, minor allele frequency, and linkage disequilibrium, and develop a method that separates these power-related confounders from biological differences. Finally, we show that analysing studies together boosts power, increasing portability. Our results provide a framework to interpret multiple eQTL studies for how genetic variation affects gene regulation across cohorts and ancestries.

## Introduction

The phenotypic effect of germline variants is often realised through gene regulatory effects [1, 2]. Population-level expression quantitative trait loci (eQTL) studies link genetic variants to inter-individual differences in stable gene expression levels [3], and have played a large role in the identification of genetic variation that underpins processes such as the development and progression of disease [4, 5]. However, these gene regulatory effects can be modulated by genetic background and macro-scale environmental variation [6]. Aside from these mechanisms, the association between a variant and expression levels can also be affected by study sample features, such as the variant’s frequency or by its correlation to other variants across the genome (linkage disequilibrium) [7, 8]. These genetic and environmental differences can vary widely across human populations [9, 10], giving rise to differing estimates of effect sizes across ancestries. Known as the “portability problem”, predictive models developed from genetic associations, such as eQTL studies or polygenic risk scores, can be up to 50% less accurate when applied out-of-sample, especially to groups that differ substantially along either environmental or genetic axes [11–13]. The portability problem remains one of the largest challenges in the equitable development of precision medicine, and, unless addressed, has the potential to exacerbate existing healthcare inequities [14–17]. Despite efforts to generate more ancestrally and environmentally diverse eQTL discovery cohorts (e.g. [18–22]), the vast majority (89%) of participants in the eQTL Catalogue and other public repositories are drawn from European genetic ancestries [23, 24], meaning the cellular consequences of genetic variation remain poorly characterised across ancestries and environments.

Past multi-ancestry studies have applied diverse definitions and methods to assess eQTL portability. For example, portability has been quantified using genetic correlations between eQTL studies [25, 26]; Bayesian tests of colocalisation between eQTL summary statistics [27, 28]; credible set overlap in fine-mapping analyses [29]; significance of individual eSNPs in both populations [21, 30, 31]; or thresholds on cross-population effect size ratios [20]. Each of these metrics describes different statistical questions about the heterogeneity of eQTL signals across cohorts, but it is not always clear how these metrics relate to each other, and whether they support consistent conclusions about the degree of effect sharing between populations. For the purposes of meta-analysis, it is currently difficult to interpret the results from different portability metrics across multiple cross-ancestry eQTL association studies to form a coherent view of the regulatory landscape across populations and ancestries.

Recent work focusing on both GWAS and eQTL mapping argues that the effects of genetic variants are largely similar across ancestries, with limited statistical evidence for “gene by gene” (GxG) interactions such as epistasis [26, 32, 33]. Instead, study size, linkage disequilibrium (LD), and minor allele frequency (MAF) appear to be the primary factors hindering portability across populations. However, portability analyses are typically restricted to a limited number of ancestries, tissues and RNA-seq technologies motivating further work to quantify how these forces drive estimates of portability across diverse cohorts. Crucially, this raises the question of whether apparent population-specificity in published eQTL studies reflects true biological heterogeneity or is largely an artefact of variable statistical power. Disentangling these two sources of non-portability therefore requires methods that can use known differences in study power — arising from MAF, sample size, and LD — to separate biologically driven population-specific effects from statistical artefacts. It also motivates statistical methods that can meta-analyse eQTL summary statistics across ancestries to obtain robust estimates of genetic effects on gene expression and more informed estimates of portability between populations.

In this work, we use published summary statistics from ten study cohorts spanning multiple tissues, genetic ancestries and RNA-sequencing technologies to systematically compare measures of eQTL portability, and develop methodologies to meta-analyse eQTL studies. First, we compare how different portability metrics used in past literature relate to each other and influence estimates of cross-ancestry eQTL sharing. Next, we quantify the major drivers of non-portability, including study size, MAF, and LD. We develop meta-analysis approaches that incorporate MAF-adjusted expectations of eQTL sharing to differentiate between non-portability due to statistical power differences (MAF and sample size) and other unmodelled population-specific effects. Finally, we compare multivariate shrinkage methods and fixed-effect meta-analysis and show how they can be used to integrate eQTL signals across ancestries, allowing for better-powered identification of population-specific eQTLs.

## Methods

### Datasets

To develop a list of candidate eQTL studies to include in our analysis, we searched for publicly-available eQTL study summary statistics in March 2023, using the eQTL Catalogue v5 [24] and Google Scholar for eQTL studies with sample size (*>* 70; Table S1). We excluded studies if they did not provide publicly-available summary statistics with sufficient information to calculate effect size and standard error for every tested SNP-gene pair (i.e., not just for independent or significant variants only).

We generated ‘sets’ of eQTL studies matched on important technical and biological factors: gene expression assay technology, tissue, and participant health status, age, and gender, excluding sets of studies from further analysis if they did not consist of at least three studies or cohorts. Additionally, we aimed to construct our Study Sets such that each set would allow us to measure cross-population effects; thus we required that each set contain at least one study where most (*>* 70%) participants came from a single, broad, non-European ancestry group. In total, we assembled three sets of matched eQTL summary statistics with specific ancestry-associated groups (Table 1). Set 1: CD14^+^ monocytes microarray gene expression data from European-American, African-American and Hispanic healthy adult donors. Set 2: whole blood RNA-seq data from Puerto Rican, Mexican-American and African-American child donors (healthy and asthmatic). Set 3: whole blood RNA-seq data from European, European-American and Indonesian healthy adult donors. Study Sets 1 and 2 are each drawn from a single publication ([25] and [29], respectively), and eQTL summary statistics within each Set were generated using a consistent methodology. In Study Set 3 the three European cohorts were sourced from the eQTL Catalogue, which enforces a consistent computational pipeline for studies published to the platform (Table 1), but the Indonesian cohort was processed separately, using a similar, but not identical pipeline [20].

**Table 1.** Sets of matched studies included in analysis.

| Summary statistics set details |  |  |  | Matched characteristics |  |  | Data source |  |
| --- | --- | --- | --- | --- | --- | --- | --- | --- |
|  | Group identifier | Cohort | n | Biological |  | Technical |  |  |
|  |  |  |  | Tissue | Cohort characteristics | Technology | eQTL cat? | Reference |
| Set 1 | European-American | MESA [36] | 578 | CD14+ Monocytes | Healthy adults (aged 45–84) | Microarray | No | [25] |
|  | Hispanic |  | 352 | CD14+ Monocytes | Healthy adults (aged 45–84) | Microarray | No | [25] |
|  | African-American |  | 233 | CD14+ Monocytes | Healthy adults (aged 45–84) | Microarray | No | [25] |
| Set 2 | Puerto Rican | GALA II [37] & SAGE [38] | 893 | Whole blood | Healthy / asthmatic children (aged 8–21) | RNA-seq | No | [29] |
|  | Mexican-American |  | 784 | Whole blood | Healthy / asthmatic children (aged 8–21) | RNA-seq | No | [29] |
|  | African-American |  | 757 | Whole blood | Healthy / asthmatic children (aged 8–21) | RNA-seq | No | [29] |
| Set 3 | European | Estonian Biobank | 471 | Whole blood | Healthy adults (aged $\geq 18$) | RNA-seq | Yes | [39] |
|  | European-American* | GTEx v8 | 379 | Whole blood | Healthy adults (aged 20–79) | RNA-seq | Yes | [40] |
|  | European | TwinsUK | 195 | Whole blood | Healthy adult females (aged 16–98) | RNA-seq | Yes | [41] |
|  | Indonesian | Natri *et al.* | 115 | Whole blood | Healthy adult males (aged 18–76) | RNA-seq | No | [20] |

\*European-American is the predominant, although not exclusive, group identifier used for GTEx v8 whole blood donors. The exact ancestry composition of whole blood donors is not publicly listed. However, per the eQTL catalogue v5, only 11% of all individuals in the GTEx v8 cohort (all tissues) have unassigned, or dissimilar ancestries to the 1000 Genomes Project EUR super-population [24]. Thus, $\geq 80\%$ of samples in this whole blood eQTL dataset are likely of European-American ancestry.

We restricted our analysis to bi-allelic SNPs, and ensured the reference and alternative al-leles matched between studies, flipping MAF and effect-size sign where necessary. Most studies implemented a MAF *>* 0.05 filter as part of their eQTL discovery pipelines, so we only retained SNPs where this requirement was met across all studies within each Study Set. Study Set 1 [25] summary statistics did not report allele frequencies for any tested SNPs, so for this Study Set, we instead filtered out variants with MAF *<* 0.05 in the gnomAD v2.1.1 [34] groups most closely related to each study population (respectively: North-Western European, excluding Finns, for the European-American cohort; Hispanic/Admixed-American for the Hispanic cohort; African/African-American for the African-American cohort). We obtained gnomAD v2.1.1 allele frequencies from the MafDb.gnomAD.r2.1.hs37d5 v3.10.0 R package [35]. As above, this approach reflects a likely real world scenario where publicly available information is incomplete. After unifying and filtering each of the Study Sets we retained 21,979,751, 48,121,383, and 20,672,191 tested SNP-gene pairs and 9,798, 13,672 and 10,788 genes in Study Sets 1, 2, and 3 respectively.

### eQTL discovery

To identify statistically significant eQTLs within each study we used two rounds of Benjamini–Hochberg (BH) correction commonly used in past studies [42]. Briefly, starting with uncorrected summary statistics we performed a first round of BH correction independently within each gene. We then extracted the adjusted p-value for the most significant SNP within each gene and performed a second round of BH correction across these SNPs. Genes where at least 1 SNP met an FDR threshold of *<* 0.05 after the second correction were deemed eGenes with a significant eQTL; for each of these genes the most significant SNP was deemed the ‘lead eSNP’. We also defined a second set of eSNPs that we refer to as ‘all eSNPs’, which were the SNPs that had nominal p-values before any FDR correction ≤ the uncorrected p-value of the least significant lead eSNP. In downstream analyses we considered the corrected statistical significance values as well as the magnitude of the association between gene expression levels and genotype (the effect size, *β̂*).

### Determining cross population portability

To determine eQTL portability at the level of individual eSNPs, we considered two metrics used in past studies. The first is *statistical significance*, where an eSNP is deemed portable if it is below a given FDR threshold in two studies or cohorts (e.g. *FDR <* 0.05 in both) [21, 30, 31]. We compared this approach to using the *effect size ratio*, whereby an eSNP is considered portable if the ratio of effect sizes, 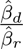, between the discovery and replication cohorts is within a given range (e.g. between 0.5 and 2). Previous work using effect size ratios has relied on effect sizes shrunk via multivariate adaptive shrinkage (MASH), without directly incorporating the standard errors of the estimates [20, 43]. We therefore compared three variants of the effect size ratio approach: *uncertainty-aware*, in which an eSNP is classified as portable only if the effect size ratio’s 95% confidence interval is within the defined boundaries; *nominal*, in which the raw effect size ratio is used directly, ignoring standard errors; and *MASH-adjusted*, in which MASH-shrunk effect sizes are used without incorporating their standard errors. In addition to comparing the overall patterns of portability across both metrics, we assessed sensitivity to threshold choice, evaluating FDR cutoffs of 0.01, 0.025, 0.05, and 0.1, and effect size ratio boundaries of 0.2, 0.5, 0.75, and 0.9.

At a gene level we also compared two approaches common in prior literature. Under the *statistical significance at the gene level* approach we deem an eGene portable from one study to a second if in both cases the gene contains a significant eSNP, regardless of whether it is the same SNP in both studies. As a second approach, we performed *colocalisation* analysis with the *coloc* v5.2.1 R package using the Approximate Bayes Factor implementation (*coloc.abf*, [44]). We called eGenes portable if the posterior probability of the two cohorts sharing a common causal variant (CCV) exceeded a given threshold. In this instance, we considered thresholds of 0.2, 0.5, 0.8 and 0.95. As recommended by the coloc manual for eQTL mapping studies, we set the prior probability of any given variant causally influencing gene expression (*p*1, *p*2) to 10*^−^*^4^. All other meta-parameters were also set to their recommended defaults [45].

The coloc.abf implementation assumes there is only one causal variant in the region being tested. Thus, we additionally considered using the coloc-SuSiE implementation, which relaxes this assumption [27, 46, 47]. This approach requires LD matrices from each population under consideration; these were not available for any of our study populations and we did not have access to raw genotype data to generate them *de novo*. Thus we used publicly available LD matrices from the 1000 Genomes Project, using the hg38 phase three release downloaded from the International Genome Sample Resource [48–50], selecting the most similar genetic ancestry to each study population (i.e., the EUR super-population LD matrix was used when the study sample was of majority European ancestry samples, AFR for African-American study samples, AMR for Puerto Rican, Mexican-American and Hispanic and EAS for Indonesia; [48, 49]). This approach led to generally poor model convergence, with the package’s diagnostic tools showing that these public LD matrices were insufficiently similar to the study populations (Figure S1), and we elected not to pursue it further.

We compared different portability metrics and thresholds using Jaccard similarity. For a pair of populations, Jaccard similarity counts the number of times two metrics agreed on whether an eSNP or eGene is portable divided by the total number of eSNPs or eGenes in the discovery cohort.

### LD-Scores

We downloaded LD-Scores for the four gnomAD super populations (AFR, AMR, EAS, NFE) from gnomAD v2.1.1 [34]. These scores were in GRCh37 coordinates, so we used UCSC lift-over with the appropriate chain files [51, 52] to convert all coordinates to GRCh38, as this is the genome build used by Study Set 2 and Study Set 3. Study Set 1 was in GRCh37 so we matched SNP ids directly across datasets. In this specific analysis we only used the matchable SNPs in each Study Set. In matching SNPs to the LD panels, we retained 20,074,822 of 21,979,751 (91%), 32,685,842 of 48,121,383 (68%) and 16,266,668 of 20,672,191 (79%) of SNP-gene pairs in Study Set 1-3.

### Predicting eQTL portability based on MAF and study sample size

To better understand the effects of MAF and sample size on eQTL portability, we formulated a method that uses the difference in MAF and sample size to predict the portability of summary statistics between study cohorts. Full derivation and simulation results can be found in the Supplementary Note. Briefly, the core assumption and motivation of this model was to assume that for any pair of matched studies, observations come from the same distribution:

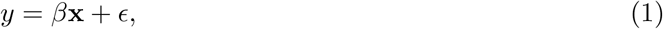

where *y* is the observed cellular phenotype (expression level for a given gene, in this case), **x** is a particular SNP, and *ϵ* is a normally distributed experimental noise term with unknown variance. Thus for two sets of observation generated from Eqn. 1, the difference in their estimators *β̂, ŝ_β̂_* would be due to sampling variation, chiefly influenced by the distribution of dosages in the **x** term (i.e., MAF) and the total number of observations in each study. To assess how well such a model encapsulates real data we conceptualised the following prediction framework:

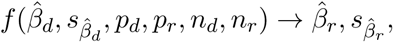

where *β̂_d_, s_β̂_*_d_ are, respectively, the estimated eQTL effect size and its variance in the discovery cohort. *p_d_, p_r_, n_d_*, and *n_r_* are the allele frequency and number of observations in the discovery cohort and replication cohort, respectively. 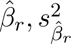 are, respectively, the predicted effect size and its variance in a repeated experiment with a different population size and/or allele frequency. With this framework we test three different ways sampling variation could impact test statistics in repeated eQTL studies:

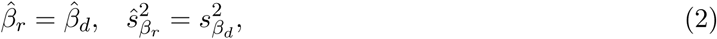

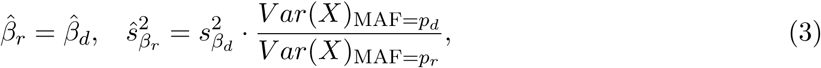

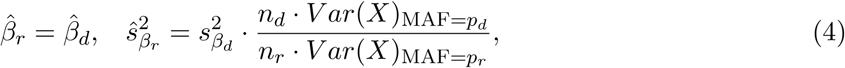

where 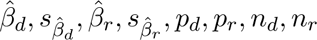 are defined as above. Var(X) is the SNP dosage variance estimated from MAF assuming HWE. Eqn. 2 simply assumes that repeated eQTL experiments should produce the same summary statistics. Eqn. 3 assumes that repeated test statistics reproduce proportionally to the difference in power due to allele frequency. Eqn. 4 assumes test statistics reproduce in repeated experiments proportional to the difference in MAF and sample size differences. We can understand these terms through the expected values of *β̂, ŝ_β̂_*

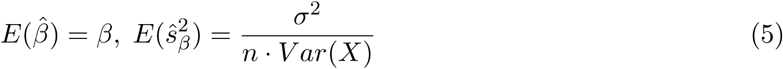

where *σ*^2^ is the variance of the residuals in Eqn. 1. We note that the expected value of *β̂* is always the true value of the data generating function Eqn. 1, whereas its expected variance depends on *σ*^2^ ∼ *ϵ*, the population size and *Var*(*X*). We note that assuming HWE *V ar*(*X*) relates to MAF (*p*) as *E*[*Var*(*X*)] = 2*p*(1 − *p*). Thus to realise our model (Eqn. 4) we assume that *β̂_d_* = *β̂_r_*, but adjust *ŝ_βr_* using Eqn. 5 assuming sample size and MAF of replication cohort as if it were the same as the discovery cohort, and unmodelled variance is equal. Thus this model gives us a framework to understand how p-statistics are influenced by MAF and population size, noting that the eQTL test statistic is equal to 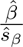.

### Meta-analysis of eQTL summary statistics studies across ancestries using adaptive shrinkage

Multivariate adaptive shrinkage (*mash*) is an empirical Bayesian method that shares signal/statistical power between repeated association studies under different conditions. The model was originally conceptualised to meta analyse summary statistics for different tissue/cell types within the same cohort of eQTL study [43]. Here, each “condition” is a different cohort within a particular Study Set. To assess the possible gain in power and cross-population portability from using *mash* to combine test statistics, we used the R package mashr v0.2.50 [43] following the best-practices provided in the vignettes ‘Accounting for correlations among measurements’ [53] and ‘eQTL analysis outline’ [54] by the package developers. The data driven covariance matrices were fitted to all principal components. We used the mashr-calculated metric, *LFSR* (local false sign rate), instead of globally corrected p-values, as our statistical significance metric as this metric already includes a high degree of multiple-test correction by virtue of the shrinkage aspect of MASH analysis. We compared MASH to fixed-effect meta-analysis which simply finds the mean effect weighted by standard errors across the pooled samples [55].

We fit mashr and fixed effect meta-analysis to all studies within each Study Set. Finally, as a means to measure the effective power gain by meta-analysing studies, we employed a leave-one-out cross-validation analysis. Here, we fit MASH and fixed effect meta-analysis to different subsets of each Study Set, leaving one study out, then compared the ability of meta-analysed studies to predict the eQTLs of the left out study. For example within Study Set 1, we adjusted the African American and Hispanic cohort summary statistics with mashr, then measured how well they predicted the European American eQTLs.

### Code and data availability

All code used in this project can be found in the following gitlab repository: https://gitlab.svi. edu.au/igr-lab/eqtl_port_fresh

## Results

### eQTL portability varies between populations and portability metrics

To investigate how different metrics of eQTL portability realise patterns of shared effects across ancestries we first collated three sets (denoted Set 1–3) of eQTL studies matched on participant characteristics, tissue, and gene expression assay technology, but spanning multiple human populations as reported by the respective study authors (Table S1). These sets span seven distinct population labels/genetic ancestries, as reported by the respective studies. Set 1 contains eQTLs identified in a single study of monocytes from healthy donors across self-reported European-American, Hispanic and African-American individuals [25]. Set 2 contains eQTLs identified in whole blood from healthy and asthmatic children of self-reported African-American, Puerto Rican and Mexican-American ethnicity [29]. Set 3 contains eQTLs identified in whole blood from four healthy adult donor cohorts: a European-American cohort [40], two distinct European cohorts [39, 41], and an Indonesian cohort [20].

Overall, the number of eSNPs and eGenes was generally correlated with the sample size of each cohort (Figure 1, Figure S2). For example, we detected the most associations in the Mexican-American ancestry cohort of Study Set 2 (the second largest across all cohorts, n = 784), with 3,369,748 eSNPs reaching nominal significance across 12,822 significant eGenes. Conversely, the Indonesian ancestry cohort of Study Set 3 had the smallest sample size (n = 115), and the fewest number of associations, with 144,063 eSNPs and 2,452 eGenes (Figure S2).

**Figure 1.**
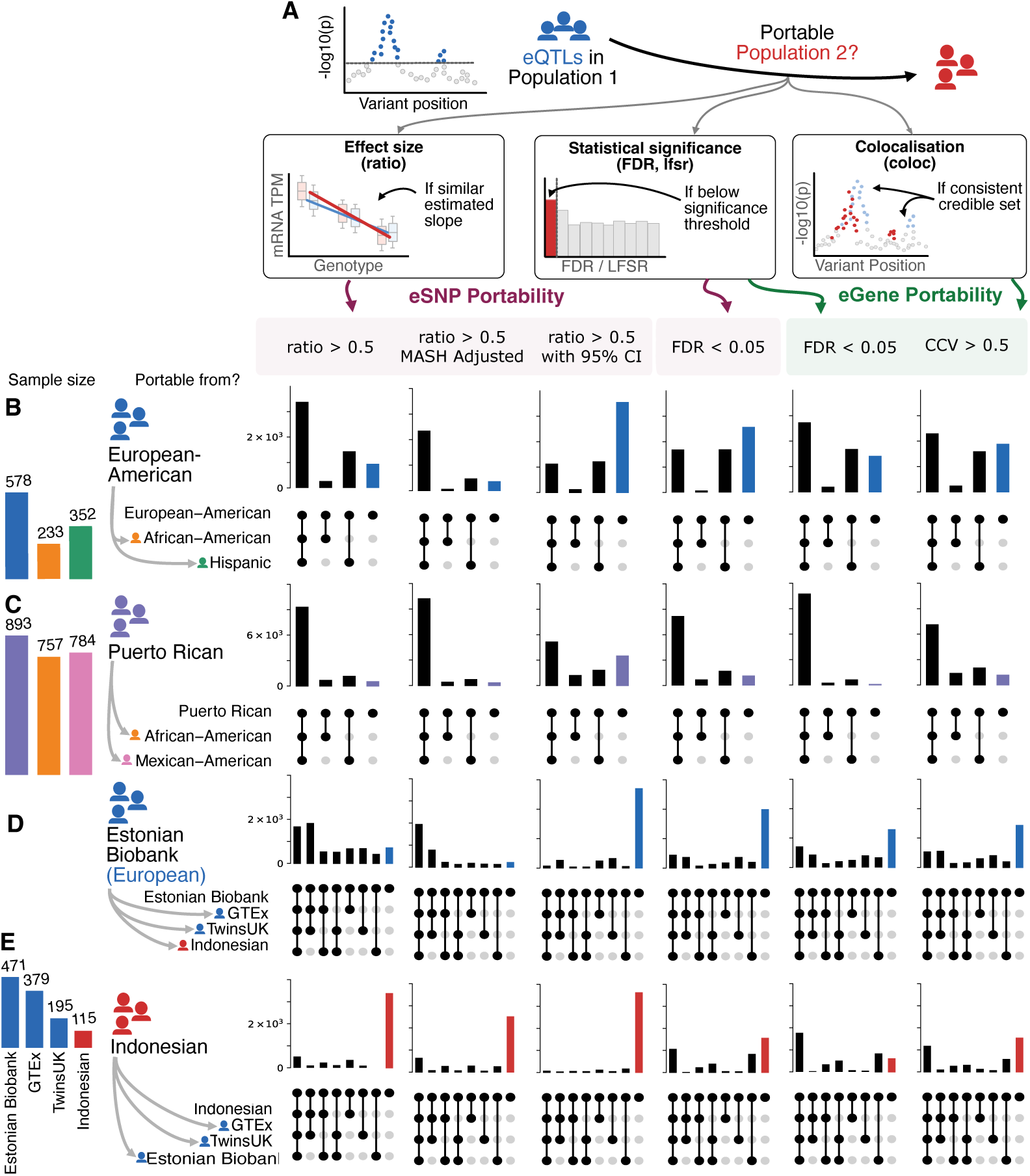
Portability of eQTLs across three metrics (significance, effect size, colocalisation) for lead eSNPs and eGenes. **(A)** Overview of the three metrics of eQTL portability tested in this study. **(B)** Portability of European-American monocyte eQTLs (4,785 eGenes / lead eSNPs) to matched African-American and Hispanic studies (Set 1). **(C)** Portability of Puerto-Rican whole blood eQTLs (11,625 eGenes / lead eSNPs) to matched African-American and Mexican-American studies (Set 2). **D.** Portability of Estonian Biobank (European) eQTLs (5,558 eGenes / lead eSNPs) to two matched European ancestry study samples, and one Indonesian study sample (Set 3). **E.** Portability of Indonesian whole blood eQTLs (2,452 eGenes / lead eSNPs) to three matched European/European-American ancestry study samples in (Set 3). **ratio** *>* **0.5** portable if the ratio of effect sizes is between 0.5 and 2. **ratio** *>* **0.5 (MASH Adjusted)** portable if the ratio of MASH adjusted effect sizes is between 0.5 and 2. **ratio 95% CI** *>* **0.5** portable if the 95% confidence interval of the SNP effect size ratio is between 0.5 and 2. **FDR** *<* **0.05** portable if the eSNP/eGene are significant in both populations. **CCV** *>* **0.5** eGene is portable in colocalisation analysis with posterior probability *>* 0.5.

Using the matched SNPs within these studies we then compared eQTL portability within each Study Set across four different metrics commonly used in past literature applicable to summary data (Methods). To do this we compared pairs of cohorts within a Study Set, defining the “discovery cohort” as the study originally used to identify eQTLs, and the “replication cohort”, as the study in which we measure their portability. eQTLs were defined as portable under: (1) *statistical significance* if an eSNP had significant p-values in both the discovery and replication cohorts; (2) *effect size ratio* if, for a significant eSNP in the discovery cohort, if the 95% confidence interval for the ratio of effect sizes, 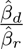, between the discovery and replication cohorts was between 0.5–2; (3) *colocalisation*, if the estimated probability of causal variant sharing (CCV) was above *>* 0.5, as calculated with the package *coloc*; (4) *statistical significance at the gene level*, whereby we define signal as portable from one cohort to another if a gene had an eQTL in both cohorts, regardless of whether signal was shared at the SNP level. For both statistical significance and effect size, we quantified portability for every eSNP (“all eSNPs”) and for the most significant eSNP in each eGene (“lead SNPs”; see Methods). Definitions 1 and 2 operate at the level of individual SNPs, whereas 3 and 4 operate at the level of genes.

Considering one cohort at a time, we measured the portability of eQTLs to all other cohorts in the same Study Set under all four definitions. Figure 1 displays patterns of sharing from one cohort to all other cohorts in the same Study Set, Figure S3-Figure S5 show comparisons across multiple thresholds. We found the proportion of eQTLs considered to be portable to at least one other matched cohort varied considerably across portability metric, and Study Set (Figure 1, Table S2). Effect ratio was more conservative than the statistical significance definition. For example in Study Set 1-3 on average 72%, 82%, and 67%, of lead eSNPs were portable to at least one other population compared to 50%,62%,32% of lead eSNPs by the statistical significance definition. For instance, the number of African-American lead eSNPs in Study Set 2 considered to be portable to the Puerto Rican cohort was 6,376 (52%), 9,008 (74%), and 8,618 (71%) for statistical significance, effect size, and colocalisation probability respectively. Additionally, the magnitude of this difference varied, even within the same set of matched studies: 79% of lead eQTLs identified in the Hispanic cohort of Study Set 1 were considered portable by the effect size ratio definition to at least one other cohort (European-American, African-American). However, by the statistical significance approach only 54% of the same eQTLs were considered portable to one of these same studies. In comparison, the difference between the proportion of eQTLs considered to be portable by effect size ratio and statistical significance approaches was much smaller for the European-American cohort in the same Study Set (portability estimates of 57% and 41%, respectively).

Past studies have also derived portability by effect size ratio without incorporating the standard errors of the effect sizes, rather using the ratio of effect sizes adjusted with multivariate adaptive shrinkage (MASH) ([20, 43]) (methods). Thus we also compared two further approaches of calling portability, first whether the nominal effect sizes were within the cutoff ratio (regardless of standard error), and also whether the MASH adjusted effect sizes were in the cutoff threshold. We found both of these approaches were more permissive at calling eQTL portability than uncertainty-aware methods. In Study Set 1-3 using an uncertainty unaware effect ratio between 0.5 and 2: 82%, 90% and 73% were portable to at least one other population, and after MASH adjustment this rose to 90%, 93% and 78%.

The exception to these trends were the Indonesian effect sizes in Study Set 3, which were poorly correlated with the other European studies in that set, such that portability was substantially higher when using a significance definition (62% of lead eSNPs portable to at least one European population) than an effect size ratio one (14% of lead eSNPs uncertainty-aware, and 31% uncertainty unaware). We speculate that the relatively small sample size of the Indonesian study (n=115, vs *n* ≥ 195 for the other cohorts in the Study Set) may explain this difference. Alternatively, small differences in the data analysis pipeline used in the Indonesian study may also mean the scale of effect size estimates are inconsistent with the rest of the studies in the Set. The Indonesian summary statistics were drawn from previous work by our team [20], while summary statistics in all other cohorts were downloaded from the eQTL Catalogue which uses a similar, but not identical, pipeline.

We more formally examined the impact of the choice of portability metric on labelling an eQTL as portable by calculating Jaccard similarities between results (Table S3). Using the uncertainty-aware effect size ratio and the statistical significance approach labelled partially overlapping sets of eQTLs as portable across cohorts within a Study Set: considering only lead eSNPs, these two measures agreed on average 80%, 77%, and 71% of the time for Study Sets 1, 2, and 3, respectively (Table S3). We also compared portability and Jaccard similarities across multiple thresholds for both SNP-level and gene-level metrics, to understand how threshold choice affected the agreement between metrics (Table S2, Table S4). Amongst all pairwise comparisons of Study Set 1 lead eSNPs, the mean Jaccard similarity between *FDR <* 0.05 and the nominal effect size ratio at cutoffs of 0.2, 0.5, 0.75, and 0.9 was 70%, 81%, 78%, and 60%, respectively. Comparing instead the uncertainty-aware effect size ratio to *FDR <* 0.05 portability, the same cutoffs (0.2, 0.5, 0.75, 0.9) agreed with *FDR <* 0.05 portability 91%, 80%, 49%, and 42% of the time. The more relaxed thresholds showed higher consistency with *FDR <* 0.05, as wide effect ratio cutoffs converge toward simply requiring effects to be nominally significant in both populations. Finally, for MASH-adjusted effect size ratios, the mean Jaccard similarity between *FDR <* 0.05 and each cutoff (0.2, 0.5, 0.75, 0.9) was 65%, 73%, 63%, and 56%, respectively. Together these results show that different portability metrics produce different patterns of portability even accounting for different stringency thresholds.

When considering portability at the gene level, we observed marked differences between the two portability definitions. Estimates of portability were consistently higher when we only asked whether a gene had at least one significant eQTL in both populations than when we formally tested for co-localisation of causal variants (Figure 1, Figure S5, Table S2). Notably, the difference here was highest in Study Set 2, which has the largest sample sizes (mean difference in the number of portable eGenes across all studies between the two methods = 23.6%). Taken together, our results indicate that some differences across metrics largely reflect stringency and discovery power, but also that in many cases, using different portability metrics can yield markedly distinct patterns of sharing regardless of the stringency of the thresholds set. Together, these results reveal how different interpretations of eQTL portability applied in past studies reveal different patterns of eQTLs across populations.

### Study size, MAF and LD structure contribute to eQTL portability

The observations above prompted us to better understand the factors that drive portability differences between studies. A major aim of many eQTL portability analyses is to enable the identification of eQTLs that by virtue of being non-portable across populations are promising candidates for uncovering gene-by-gene and gene-by-environment interactions. However, statistical factors unrelated to these processes can confound the discovery of these non-portable eQTLs and thus any subsequent investigations into context-dependent gene regulation. Since we uncovered considerable variation in the number and pattern of non-portable eQTLs across our chosen portability metrics, we were interested in exploring the technical and biological factors underlying these differences, and the extent to which statistical discovery power distorts assessments of true cross-population portability.

At a study level we found that the sample size of the replication cohort was strongly correlated, on the log scale, with eQTL portability (Figure S6), a relationship that was apparent regardless of the size of the initial discovery cohort. The relationship was strongest when using lead SNP statistical significance as a portability metric, where it reached an *r*^2^ of 0.66 for statistical significance and 0.68 for uncertainty-aware effect ratio. This relationship weakened when considering all SNPs (*r*^2^ = 0.31), or effect size ratio definitions of portability that do not include standard errors (*r*^2^ = 0.31 for lead eSNPs, *r*^2^ = 0.16 for all eSNPs). We also observed that, as was the case with eSNPs, the association between the proportion of portable eGenes and log study size was consistently strong (gene-wise statistical significance *r*^2^ = 0.70; colocalisation *r*^2^ = 0.62).

Next, we examined the relationship between portability and three other known SNP-level contributors to portability: i) effect size (*β̂*), ii) minor allele frequency (MAF), and iii) linkage disequilibrium (LD). We measured the association between each of these features and our two SNP-wise metrics of portability as well as the difference in log-transformed p-value between the discovery and the replication cohort for any given eSNP, which we use as a superficial measurement of the difference in statistical support for an eQTL across cohorts (Figure S7-Figure S12), computing these associations for both the set of lead eSNPs and for all eSNPs. Note that when calculating effect size ratio, we measured correlation to the absolute value of the log ratio, i.e. 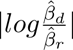. There was a small number of eSNPs that had different directions of effects in cohort pairs, which confounded the log transform. We discarded these SNPs to avoid negative effect size ratios to keep the metric on a continuous scale and simplify interpretation. Across each possible pair of populations this resulted in an average of 5.0%, 7.6% and 8.0% of all eSNPs being removed for the effect size ratio analysis specifically.

Across all Study Sets and portability metrics, eQTLs with a large absolute test statistic or effect size in the discovery cohort were more likely to be portable to the replication cohort, as larger effects are easier to detect, even taking into account effect size inflation driven by Winner’s Curse [42]. Unsurprisingly, test statistic in the discovery cohort therefore was the strongest SNP-level driver of portability that we identified, with consistently high correlations between *β̂* and portability across all 3 Study Sets (Figure S7-Figure S12). For example, for the lead eSNPs in each population, the Spearman *ρ* between eSNP p-value in the discovery cohort and the p-value in replication cohort across all 3 Studies was at minimum 0.44 going from the Indonesian cohort to the GTEx cohort, to at maximum 0.88 from the Mexican cohort to the Puerto Rican cohort in Study Set 2. (Figure S7). Similar ranges were also observed considering the set of all eSNPs, as well as when using the effect size ratio portability metric We then examined the relationship between MAF and portability (Figure 2, Figure S7-Figure S12). We found that MAF in the replication cohort was consistently associated with statistical significance, especially when considering lead eSNPs (mean *ρ*: Set 1 = −0.13 ± 0.045; Set 2 = −0.18 ± 0.037; Set 3 = −0.13 ± 0.028; Figure S7); the set of all eSNPs showed similar but weaker levels of association (mean *ρ*: Set 1 = −0.11 ± 0.050; Set 2 = −0.13 ± 0.045; Set 3 = −0.11 ± 0.020; Figure S10). The association with MAF was also strong when we considered differences in log p-value between cohorts (mean *ρ* across lead eSNPs: Set 1 = 0.20 ± 0.062; Set 2 = 0.37 ± 0.030; Set 3 = 0.091 ± 0.019; Figure S8). However, effect size ratio between cohort pairs showed lower levels of association with replication cohort MAF for both the set of lead eSNPs (mean *ρ*: Set 1 = 0.05 ± 0.052; Set 2 = 0.065 ± 0.028; Set 3 = 0.041 ± 0.0097; Figure S9) and set of all eSNPs (mean *ρ*: Set 1 = 0.0061 ± 0.047; Set 2 = 0.037 ± 0.045; Set 3 = −0.019 ± 0.014; Figure S12). These results demonstrate that eQTLs considered to be non-portable by effect size ratios or statistical significance typically have lower MAF in the replication cohort than portable eQTLs; consistent with the hypothesis that differences in study power hinder the identification of truly non-portable eQTLs.

**Figure 2.**
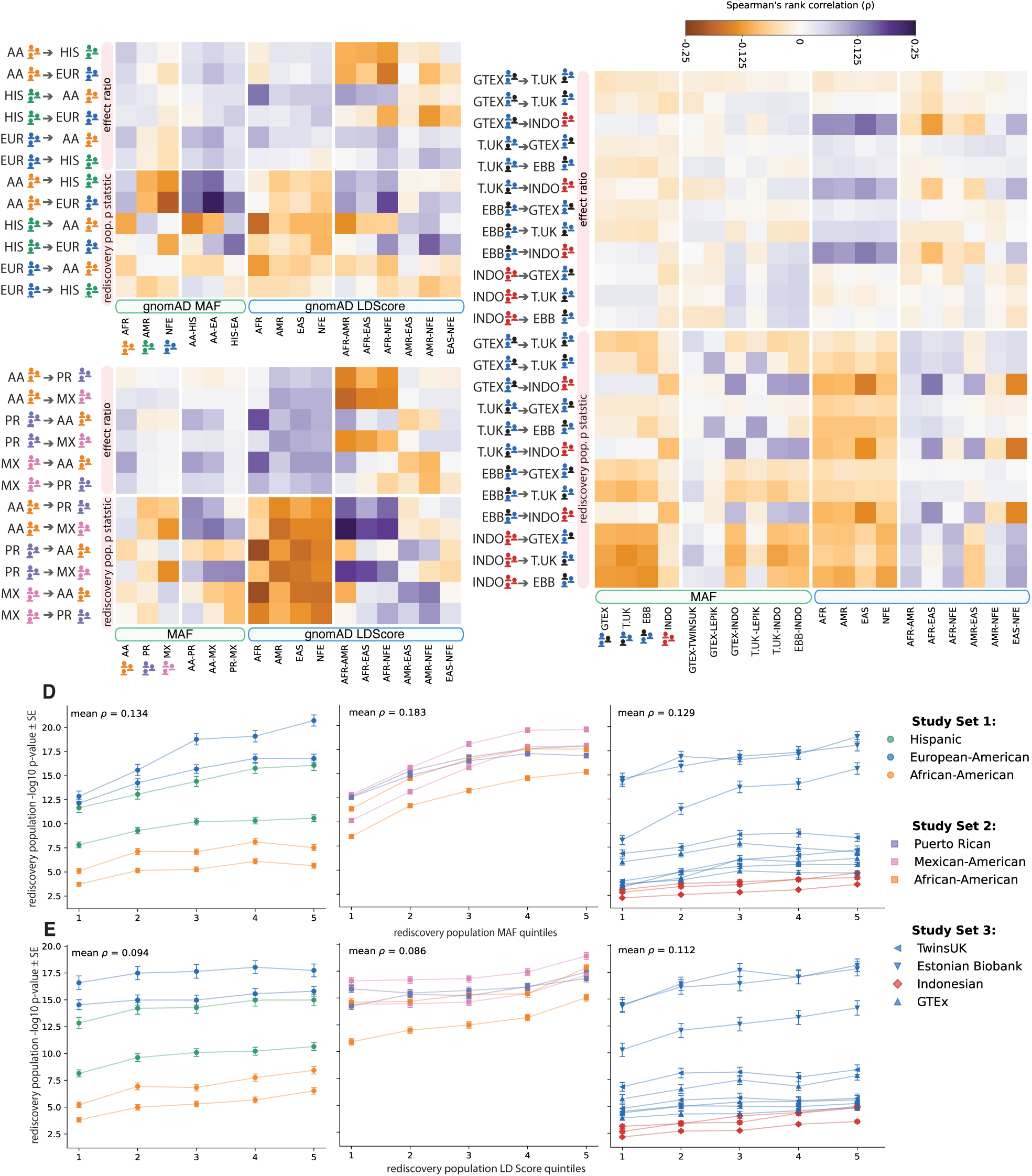
MAF and LD differences between cohorts impact eSNP portability. **(A-C)** Spearman Correlation between SNP-wise portability metrics (nominal p-value in replication study, effect size ratio) and SNP-level contributors to portability (MAF and gnomad LD-Score) in Study Set 1-3 (A-C respectively) among discovery cohort lead eSNPs. **(D)** the mean − log 10 p-value of replication cohort lead eSNPs stratified into 5 bins based on replication cohort MAF. **(E)** the mean − log 10 p-value of replication cohort lead eSNPs stratified into 5 bins based on replication cohort LD-Score. Results pertain to the set of all significant lead eSNPs. See Figure S10-Figure S12 for other data subsets. **AFR:** African; **AMR:** Admixed American; **EAS:** East Asian; **NFE:** Non-Finnish European.

In general, the correlation between MAF and population size was proportional to the replication study size, especially considering the relationship between log p-value and differences in MAF. For example, when considering portability of lead eSNPs to the Mexican-American cohort of Study Set 2 (*n* = 784) the association between MAF and p-value was on average *ρ* = −0.22, but when considering replication in the Indonesian cohort (*n* = 115) this value drops to *ρ* = −0.13 Figure S7. Indeed, regressing the size of the replication cohort on the correlation between replication cohort MAF and replication cohort p-value had a linear relationship among lead eSNPs (*r*^2^ = 0.24; Figure S13); this association is significantly strengthened when we consider the relationship between the difference in MAF and the difference in log p-value between the replication and discovery cohorts (*r*^2^ = 0.65; Figure S13A). This phenomenon also persisted in considering all eSNPs to a weaker extent (Figure S13B).

Finally, we considered the role of LD in assessing portability. Given our reliance on public summary statistics without associated in-sample LD matrices available, we used LD-Scores from multiple gnomAD super-populations (AFR, AMR, EAS, NFE) as our LD metric. This decision was motivated by the fact that cohorts including minoritised populations tend not to be large enough to construct reliable LD maps. Our use of reference LD instead of in-sample LD thus reflects typical challenges encountered by researchers examining eQTL portability. LD-Score is a cumulative measure of correlation between genotype at the focal locus and all other variable sites genome-wide. Since eQTL mapping uncovers statistical associations between an eSNP and eGene pair, we reasoned that LD differences between populations could explain a substantial fraction of non-portable eQTLs, as differences in haplotype structure disrupt statistical associations when considering portability even in the presence of shared biological causality, especially when considering all significant eSNPs. Indeed, LD-Score was associated with portability across both lead eSNPs and all eSNPs (Figure 2). When considering only lead eSNPs, this association was strongest for Study Set 3, the set with the smallest sample sizes (Figure S7, Figure S9). Notably in this Study Set, LD-Score was much more informative when attempting to port results from European cohorts to the Indonesian cohort than when porting them between European cohorts, likely because LD structure does not differ as much between the three European cohorts and thus cannot provide additional information (Figure 2C). For instance, in both Set 1 and Set 2, LD-Score in gnomAD African (AFR) samples was strongly predictive of statistical significance in African-American cohorts. Comparisons involving the Indonesian cohort in Set 3 revealed a much stronger association between LD-Score in gnomAD East Asian (EAS) samples than non-Finnish Europeans (NFE). This result is consistent with results from the first publication on this sample: PCA of genotype data from the Indonesian samples in the original publication revealed a close genetic similarity to 1KG-CHB samples [28], which were used by gnomAD to group samples into the EAS cluster [34] from which the LD-Score matrix was derived.

We observed that differences in LD-Score between populations can also be informative in assessing portability. For instance, when considering all eSNPs, LD-Score differences between AFR and any of the other three LD-Score matrices were negatively correlated with portability from African-American cohorts to other studies in both Set 1 and Set 2 (AA to elsewhere Set 1 mean LD-Score to p value *ρ* = −0.106, Set 2 = −0.175, Figure 2A-B), regardless of how we define it. Additionally, in Set 2, LD-Score comparisons between AMR and NFE or EAS were also strongly correlated with portability from Mexican-Americans to Puerto Ricans, and from Puerto Ricans to either Mexican-Americans or African-Americans. These patterns are surprising because genetic ancestry analyses performed in the original publication indicate that while both the Puerto Rican and Mexican-American cohorts are admixed between European-like, African-like and Indigenous American-like genetic ancestries [29], the relative proportions of these three vary substantially across the two groups. Taken together, our observations suggest that linkage disequilibrium is a substantial driver of non-portability of eQTLs. Furthermore, while all LD metrics we considered were derived out-of-sample, their ability to capture some degree of relevant regional structure suggests they could be useful in future research aiming to assess portability in an LD-aware manner.

### Correcting for MAF and population size in estimates of eQTL portability

Building on the observation that both sample size and allele frequency largely impact eQTL portability, we wanted to explicitly account for uncertainty when calling eQTLs “non-portable”. We developed a framework that assumes effects sizes are persistent between populations, but uses the difference in MAF and sample size to ascertain if a replication cohort has sufficient statistical power to detect eQTLs from the discovery cohort. This framework, inspired by [56] effectively down-samples the discovery cohort at the summary statistic level to have the same MAF and sample size as the replication cohort, allowing us to scrutinise eQTLs for portability only if they are significant in the discovery cohort after matching the sample size to the replication cohort (methods). We validated this method with simulations (Supplementary Note), and here present how well this interpretation explained the portability between cohorts in our Study Sets. We compared three possible relationships between the discovery cohort and replication cohorts test statistic, shown in columns 1–3 of Figure 3 (see Table S5 for results for all pairwise comparisons). The first (Eqn. 2) was the identity transform, which assumes that the replication study should perfectly reproduce the discovery study. The second (Eqn. 3) adjusts the summary statistics of the discovery study based on the difference in MAF of the discovery and replication cohort. The third (Eqn. 4) adjusts the test statistics of the discovery study based on both the difference in MAF and population size.

**Figure 3.**
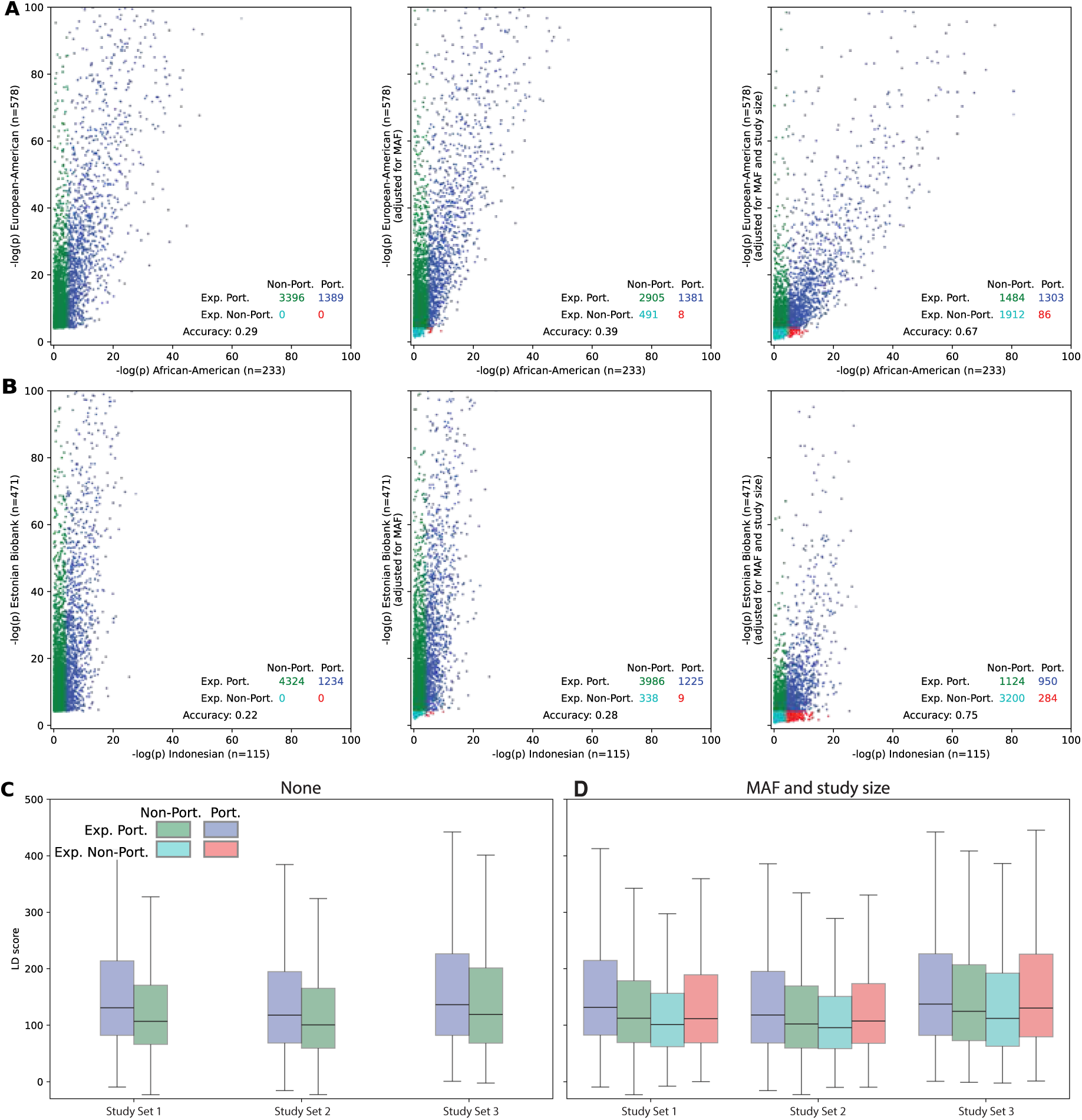
Predicted lead eSNP portability accounting for MAF and sample size. **(A-B)** Shows portability from two pairs of populations from Study Set 1 and Study Set 3 respectively. Each dot is a lead eSNP identified in the discovery cohort. The *x* and *y* axes show − log(p-value) in the replication and discovery cohorts, respectively. Columns correspond to different y-value transformations: *left* (unadjusted); *middle* (MAF-adjusted, Eqn. 3); *right* (MAF- and sample-size–adjusted, Eqn. 4). **(C)** Mean replication cohort LD-Scores for discovery cohort lead eSNPs **(D)** Mean replication cohort LD-Scores for discovery cohort lead eSNPs after applying MAF-and sample-size–adjustment (Eqn. 4). Per panel, confusion-matrix counts report observed (**Port.**, **Non-Port.**) and predicted (**Exp. Port.**, **Exp. Non-Port.**) counts of portable and non-portable eSNPs (*y* → *x* population), respectively. **Accuracy**: proportion of correctly classified eSNPs.

The outcome of these three transformations is presented in Figure 3, with columns showing the transformations associated with Eqn 2–4 from left to right. Using Study Set 1 as an example (Figure 3A), the first column presents the same results as shown in Figures 1, but at the level of individual eSNPs, because in Eqn. 2 we assume the replication study should perfectly reproduce the discovery study. As we found earlier 29% of lead eSNPs from the European-American cohort are also significant in the African-American cohort. In the middle column, we scale our expectations of portability by Eqn. 3, adjusting the test statistic of the discovery cohort as if it had the same MAF as the replication cohort. This scaling typically shrinks test statistics in the discovery cohort (European-American in this case) with low MAF in the replication cohort, such that we predict fewer eQTLs will be statistically significant in the replication cohort. In Study Set 1, our approach suggests that 491 (14%) of eSNPs initially considered “non-portable” from the European-American to the African-American cohort can be explained solely by power differences due to MAF.

Finally, in the third column we explicitly account for both MAF and sample size differences. Continuing the example of Study Set 1, the European-American cohort (*n* = 578) was much larger than the African-American (*n* = 233). Thus, our transformation applies additional shrinkage to the European-American test statistics proportional to this power difference. This results in up to 1,912 (56%) of significant lead eSNPs being shrunk outside the significance threshold and thus becoming non-portable. In other words, the observed *β̂* in the European-American cohort cannot produce a significant result under the sample size and MAF of the African-American population even in the presence of a shared biological effect. Thus, even though these eSNPs are classified as “non-portable” when we naively compare statistical significance between studies, we could never have expected them to be portable *a priori* given the power difference between the two studies. In comparison, only 6% of the “portable” eSNPs from European-American to the African-American cohorts of Study Set 1 had “non-portable” p-values after the MAF and study size transformation. Overall, accounting for power differences between the two studies allowed our approach to predict eQTL portability from the European-American to the African-American cohort with 67% accuracy (Figure 3A).

We observed a similar pattern across all other Study Set and cohort combinations. Unsurprisingly, when the sample size in both cohorts was large, the effect of transforming based on MAF and sample size was more modest. For example, in Study Set 2 after accounting for the power difference due to MAF and sample size, 81% of eSNPs were correctly predicted to be portable/non-portable from the Puerto Rican to the African-American cohort (Table S5). Comparatively, in Study Set 3, most non-portability was attributable to differences in study size; only 22% of the eSNPs in the Estonian Biobank were deemed portable to the Indonesian cohort in the analyses above. Accounting for sample size and MAF, portability from the Estonian Biobank to Indonesia could be predicted in 75% of cases *a priori* (Figure 3B). Indeed, across the three Study Sets, we were able to correctly predict if an eQTL was portable on average 73%, 77% and 61 % of the time, respectively (Table S5).

We hypothesised that LD differences might be a primary driver of non-portable eQTLs that are well powered in both populations. Before adjusting summary statistics, non-portable eQTLs had significantly higher LDScores (TP vs FP mean difference: 34.39, 25.41, and 17.94 in Study Sets 1–3 respectively, all *p <* 10*^−^*^56^; Figure 3C). After adjusting for MAF and differences in sample size, well powered portable eQTLs (TP) continued to show higher LD-Score than non-portable eQTLs (FP) across all Study Sets (mean difference: 28.22, 23.42, and 12.93 in Study Sets 1–3 respectively, all *p <* 10*^−^*^23^, and portable eQTLs also had substantially higher LD than true negatives (TP vs TN mean difference: 45.24, 37.44, and 25.43 in Study Sets 1–3 respectively, all*p <* 10*^−^*^35^. Among eQTLs that did not have the expected power to be portable, those that were portable (FP) had higher LD than those that were not (TN) (mean difference: 17.02, 14.02, and 12.50 in Study Sets 1–3 respectively, all *p <* 10*^−^*^4^. Thus these results are consistent with SNPs with high LD being able to tag more signal, thus boosting associations and portability of eQTL signals, and show how this contributed to non-portability even when accounting for differences in statistical power.

### Meta-analysed summary statistics are more portable to out-of-sample cohorts

Given that we have shown how differences in statistical power explain a large portion of non-portability, we reasoned that combining eQTL studies to improve power should better isolate biologically-driven, cohort-specific eQTLs. As power increases, the remaining non-portable eSNPs should represent a smaller and more informative set of candidates for population-specific regulation, gene-by-environment interaction, or other context-dependent effects. We therefore further explored the multivariate adaptive shrinkage (MASH) method [43], implemented in the *mashr* R package — an empirical Bayes approach that learns a prior distribution of effect sizes and their covariance across groups, then uses this prior to compute posterior estimates for all association tests. We note that in earlier sections comparing portability metrics, we showed the ratio of MASH-adjusted effect sizes to be the most permissive measure of eQTL portability used in past literature [20]. Here we further explore the potential of the MASH framework to improve power, and compare its potential and applicability to multi-cohort eQTL studies.

As expected, cohorts with larger sample sizes identified more eQTLs and eGenes both before and after MASH. Applying MASH to all populations within each Study Set significantly increased the number of discovered eSNPs by an average of 125%, 123%, and 171% in Studies 1, 2, and 3, respectively. The mean increase in eGenes was 66%, 11% and 95% in Study Set 1-3 respectively. The relative gain in power was most substantial in studies with smaller sample sizes (Figure 4A). For instance, in Set 1, the number of eQTLs detected in each study population (European-American, Hispanic, African-American) was 602,640, 332,528, 118,610 prior to, and 895,111, 809,737, and 666,128 after applying MASH analysis by the same FDR or LFSR *<* 0.05 threshold. This increased power from information sharing allowed us to detect eSNPs with lower MAF, with a consistent decrease in the minimum MAF of significant eSNPs per eGene across all studies (sign test across all 10 cohorts *p* = 9.8 × 10*^−^*^4^). This result indicates that MASH helps detect associations which were originally not statistically significant in individual cohorts due to limited observations.

**Figure 4.**
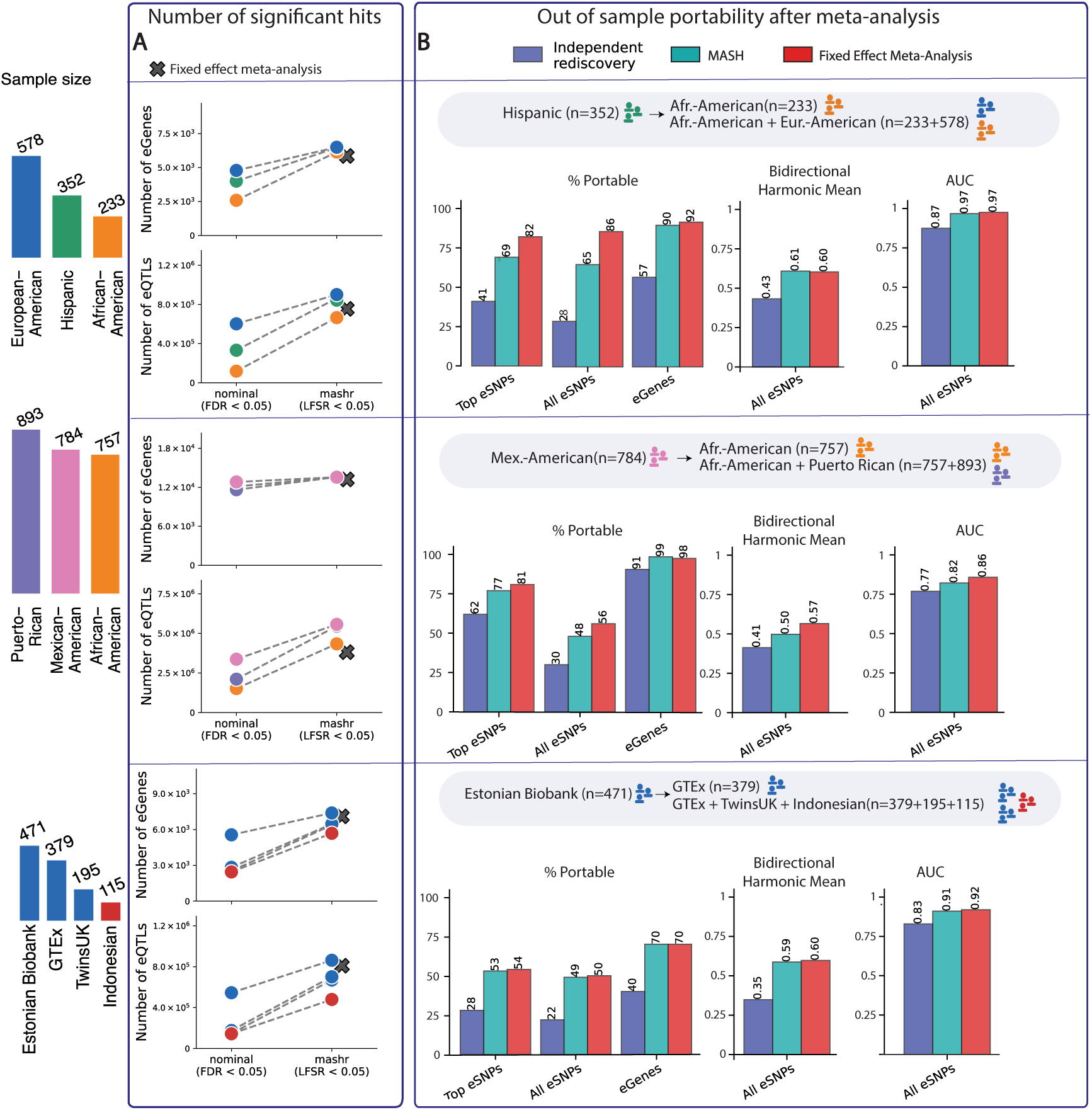
Meta-analysis of eQTL studies reduces the confounding effect of power differences across studies on portability analyses. (A) The number of eSNPs per study before and after mashr or fixed-effect meta-analysis was applied across all cohorts. (B) Comparison of portability metrics between a discovery cohort and a single replication cohort versus a meta-analysed replication cohort. **Independent replication** (dark blue) bars show portability metrics from the discovery cohort directly to the replication cohort (e.g. Hispanic to African American in the first row). **MASH** (teal) bars show the same metrics, but with the replication cohort adjusted using MASH together with the other populations in the study. **Meta-analysis** (red) bars show portability metrics from the discovery cohort to all other cohorts meta-analysed together using a fixed-effect meta-analysis. **% portable:** the percentage of eSNPs that are significant in both the discovery and replication/meta-analysed cohort. **Bidirectional harmonic mean:** the harmonic mean of the proportion of eSNPs portable from the discovery cohort to the replication/meta-analysed cohort, and the proportion portable in the reverse direction. **AUC:** the area under the receiver operating characteristic curve for the number of portable eQTLs from the discovery cohort to the replication/meta-analysed cohort, across different significance thresholds used to define portability. Note that not all metrics can be applied to all sets of eSNPs; for example, the bidirectional harmonic mean cannot be applied to top eSNPs, as lead eSNPs form different sets of SNPs across populations.

We also compared MASH to a simpler fixed-effect meta-analysis, which also increased the mean number of discovered eSNPs to a comparable degree of 115%, 64%, and 220%, and the mean number of eGenes by 54%, 8%, and 112%, in Studies 1, 2, and 3 respectively. However, unlike MASH this method simply pools all populations together losing resolution among the pooled samples.

To further validate MASH in the context of portability, we performed leave-one-out cross-validation for each Study Set. In each case one cohort was held out as the discovery set of eQTLs, and we tested how well the remaining cohorts recovered its eSNPs — either individually or after combining them by MASH or fixed-effect meta-analysis. A genuine increase in power should recover more of the held-out cohort’s eSNPs. In Study Set 1, treating the Hispanic cohort as the discovery cohort, the proportion of its eSNPs recovered by the African-American cohort was 29% (41% for lead SNPs). After MASH-adjusting the African-American cohort with the European-American cohort, this replication rose to 65% (69%). Pooling the African-American and European-American cohorts with a fixed-effect meta-analysis further increased the replication to 86% (82%) (Figure 4B). Fixed-effect meta-analysis recovered the most eSNPs; however, unlike MASH, it simply pools all populations together, losing resolution among the pooled samples. Comparisons across the other Study Sets showed similar patterns of increasing replication (Table S6–Table S8).

Recovering a larger fraction of the held-out eSNPs is not by itself conclusive. Because MASH and fixed-effect meta-analysis enlarge the significant set of the replication cohorts, the same gain would be expected simply from adopting a less conservative discovery threshold. To separate a real improvement from a mere relaxation of stringency, we summarised the same recovery across all replication LFSR thresholds at once using the AUC (Figure 4B). As a complementary bidirectional summary, we also took the harmonic mean of the portability proportions in both directions (i.e. discovery-to-replication and replication-to-discovery); this rises only when shared signal is detected consistently rather than asymmetrically. Both MASH and meta-analysis increased this AUC across every leave-one-out combination. In Study Set 1, the AUC with which the replication data recovered the held-out Hispanic eSNPs rose from 0.874 (0.888 for lead SNPs) for the African-American cohort independently, to 0.967 (0.945) after MASH-adjustment with the European-American cohort, and to 0.975 (0.947) under meta-analysis; the corresponding bidirectional harmonic mean rose from 0.43 to 0.61 (MASH) and 0.60 (meta-analysis). In Study Set 2, holding out the Mexican-American cohort, the AUC rose from 0.769 (0.886) for the African-American cohort independently, to 0.822 (0.907) after MASH-adjustment with Puerto-Rican, and to 0.858 (0.922) under meta-analysis, with the harmonic mean increasing from 0.41 to 0.50 (MASH) and 0.57 (meta-analysis). In Study Set 3, holding out the Estonian BioBank cohort, the proportion of its eSNPs recovered rose from 22% (28% for lead SNPs) for GTEx independently, to 49% (53%) after MASH-adjustment within the pooled cohorts (GTEx, TwinsUK, Indonesian), and to 50% (54%) under meta-analysis; the AUC rose in step from 0.829 (0.824) to 0.910 (0.891) to 0.919 (0.899), and the harmonic mean from 0.35 to 0.59 (MASH) and 0.60 (meta-analysis). In Study Set 3 we also tested holding out the Indonesian cohort. This showed the AUC rise from 0.902 (0.872 for lead eSNPS) for the Estonian BioBank independently to 0.915 (0.885) after MASH-adjusting the Estonian BioBank with other European-ancestry cohorts and 0.914 (0.883) under meta-analysis of all European-ancestry cohorts in the Study Set. However here the harmonic mean fell slightly, from 0.27 to 0.21 (MASH) and 0.23 (meta-analysis). This reflects the very small size of the Indonesian cohort (n=115): the backward portability of the meta-analysed cohorts (GTEx n=379, TwinsUK n=195, Estonian BioBank n=471) to Indonesian was low, at only 13% and 14% for MASH and fixed-effect meta-analysis respectively (Figure 4B).

## Discussion

Understanding patterns of eQTL sharing between genetic ancestries gives insight into the cellular basis of important diversity between populations, including the development and progression of disease [57–59]. Past literature has used diverse methods to categorise eQTLs as population-specific, which hinders comparison between studies and estimates of the true cross-population portability and replication rate. Moreover, separating power-related (i.e., MAF and study size) reasons for non-portability from other factors (such as GxE) remains under-explored in existing work. In this study we collected eQTL summary statistics spanning ten studies across multiple genetic ancestries, two tissues, and two transcriptomic technologies. We show that popular methods of computing portability reveal different levels of eQTL sharing between populations. We next demonstrate that MAF and study size are the major technical drivers of non-portability between studies, and present a method to predict if eQTLs are non-portable due to insufficient power, or other unmodelled factors. Finally, we show how multivariate adaptive shrinkage can reconcile differences between studies, and can be used in meta-analysis for deeper insight of the genetic effects on gene expression across ancestries, and reveal a smaller but better powered set of population-specific eQTLs.

We found that different metrics of eQTL portability commonly used in prior literature produce different degrees of eQTL sharing between ancestries, a result that explains how diverse conclusions about eQTL sharing between populations have been reached in the literature. For example, statistical significance is more conservative than using the MASH adjusted ratio of effect sizes. Similarly, overlap of lead eSNPs is more conservative than colocalisation or overlap of eGenes. These results partially explain differences in eSNP sharing across ancestries reported in prior studies. For example, Natri et al. conducted an eQTL study with participants recruited from Indonesian populations [20] and reported that 38% of genes show no evidence of colocalisation to GTEx [40], whereas using a similar sample size, but using statistical significance rather than colocalisation, Kelly et al. [21] found that only 17% of eQTLs from a similarly sized East African cohort were portable to GTEx. Without directly examining the raw summary statistics, there is limited scope for interpretation when comparing the two results. Furthermore, while some portability metrics were more stringent, the different patterns of portability between metrics could not be fully reconciled by adjusting threshold parameters and thus render comparisons of portability estimates between studies challenging. Our results therefore underscore the importance of sharing summary statistics from eQTL studies as the only reliable way to infer patterns of the genetic determinants of gene expression across populations by combining multiple studies.

We demonstrate that population size and minor allele frequency are associated with eQTL portability. In line with past studies, we found a strong correlation between the size of the replication sample and eQTL portability across every measure of portability tested. This correlation was weakest for effect size ratio, which could be due to the fact that lower powered tests of association (stemming from sample size and MAF) will tend to have greater variance in effect size estimates. Comparatively, estimation uncertainty is directly accounted for in the calculation of p-values and colocalisation analyses. At the SNP level we found MAF to be strongly correlated with the statistical significance definition of portability. We found this association of MAF to be stronger in cohorts with larger sample size—particularly considering the set of lead eSNP. Our results illustrate that higher sample sizes are required to obtain sufficiently precise effect size estimates and standard errors for cross-cohort comparison, enabling us to attribute residual non-portability to mechanisms beyond sampling variance, including locus-specific architecture and environmental modulation.

We have presented a simple approach for comparing eQTL portability across cohorts and study sizes, building on the observation that differences in power between studies confound efforts to identify biologically meaningful explanations for non-portability. In particular, it can be hard to determine whether a SNP is non-portable between two populations because of MAF/study size differences or because of study-specific effects such as GxE. Our approach establishes statistical boundaries when measuring portability from an arbitrary discovery cohort to a replication cohort—especially when the discovery cohort has a sample size greater than or equal to the replication cohort—by rescaling the standard errors of the discovery cohort’s slope estimates as if it had the same number of observations and MAF as the replication cohort. This approach typically down-weights the apparent significance of findings in discovery cohort proportionately to statistical power in the replication cohort. We show that this technique enables separation of SNPs expected to be portable from those expected to be non-portable due to insufficient power in the second population. We found that well powered non-portable eQTLs were depleted for LD compared to the portable eQTLs. SNPs with high LD tag more variation so even after adjusting for MAF and sample size differences LD remains a confounder that should be taken into consideration in defining biologically-driven cohort-specific eQTLs. Together these results provide a framework for identifying the SNPs most likely to be enriched for genuine study-specific environments (GxE) or ancestry-associated genetic backgrounds (GxG). Where prior work has sometimes downsampled larger cohorts to equalise power across cohorts our approach provides a more statistically principled comparison across unequal study sizes while also accounting for MAF-driven power differences, applicable to summary statistics. The high fraction of eQTLs correctly predicted as portable/non-portable using only MAF and sample size supports the view that eQTL effects are largely consistent across ancestries [26, 33]. More broadly, these results are consistent with the hypothesis that ancestry-specific genetic effects on complex traits are rare [60, 61], although generation of new datasets from diverse environments may shift these estimates in the future.

Finally, we compared standard fixed effect meta-analysis to MASH (implemented in the mashr package [43]) as means to meta-analyse eQTL summary statistics across multiple studies and ancestries, to gain more robust estimates of effect sizes and their specificity to particular populations. While MASH is widely used to model effect sharing across cell types within a study, here we demonstrate its utility for quantifying sharing of eQTL effects across ancestries within the same cell type. Adjusting effect size estimates according to learned patterns of sharing increased the number of significant eQTLs in each population and increased their portability between populations. To evaluate its generalisability, we performed a leave-one-out analysis in which one population was excluded from fitting and then used solely for evaluation. MASH yielded more robust predictions in the held-out population, as reflected by greater replication/portability (e.g., increased AUC across FDR thresholds). Comparing MASH to fixed effect meta-analysis, both methods produced similar portability to held-out samples under the harmonic mean and AUC metrics, however mashr still simultaneously keeps per-population estimates while fixed effect meta-analysis reduces this resolution. MASH did slightly outperform the fixed effect model in Study Set 3, which had 4 cohorts processed with two different pipelines (compared to 3 in Study Sets 1-2, always uniformly processed), suggesting that the MASH approach might scale well as the number of populations in the meta-analysis increases. These gains are consistent with improved power from sharing information across cohorts. We therefore recommend applying mash when comparing eQTL summary statistics across ancestries to obtain more accurate estimates of SNP effects on gene expression. We also note that eSNPs remaining non-portable after MASH are natural candidates for enrichment of population-specific signals and interactions with study-specific environments (GxE) or genetic backgrounds (GxG), in line with prior observations [20].

## Acknowledgements

This work was supported by NHMRC Ideas Grant 2020501 to IGR and CBDA and NHMRC Investigator Grant 1195595 to DM. St Vincent’s Institute acknowledges the infrastructure support it receives from the National Health and Medical Research Council Independent Research Institutes Infrastructure Support Program and from the Victorian Government through its Operational Infrastructure Support Program. The funders had no role in study design, data collection and analysis, decision to publish, or preparation of the manuscript.

## Contributions

PMG: Conceptualisation, Methodology, Software, Formal analysis, Investigation, Data Curation, Writing - Original Draft, Writing - Review & Editing, Visualisation

IJB: Conceptualisation, Methodology, Validation, Formal analysis, Investigation, Data Curation, Writing - Original Draft, Writing - Review & Editing, Visualisation.

CBDA: Conceptualisation, Supervision, Writing - Review & Editing, Funding acquisition

DJM: Methodology, Writing - Original Draft, Writing - Review & Editing, Supervision, Funding acquisition

IGR: Conceptualisation, Methodology, Writing - Original Draft, Writing - Review & Editing, Supervi-sion, Funding acquisition

## Supplementary Note

To formulate the relationship between minor allele frequency (MAF), sample size (*n*) and eQTL portability, we assumed that genetic variants always have the same effect on gene expression regardless of experiment so any difference in portability was due solely to statistical factors.

We thus created a model mapping how sample size and allele frequency affect QTL portability in a repeat experiment. Specifically, we define a function:

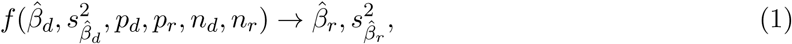

where 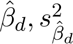 are the estimates of QTL effect size and its variance in the focal (‘discovery’) sample. *p_d_, p_r_, n_d_,* and *n_r_* are the allele frequency and number of observations in the focal experiment and repeat experiment, respectively. *β̂_r_* and 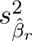 are the predicted effect size and its variance in a repeated experiment with a different sample size and/or allele frequency.

Let *X* be a random variable denoting the genotype dosage at a given SNP. We assume that all SNPs are in Hardy-Weinberg Equilibrium (HWE) and that the SNP genotypes are encoded as allele counts (i.e., dosage), thus a value in {0, 1, 2}. Accordingly, we can define the following dosage probabilities:

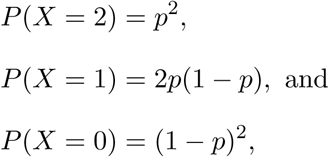

where *P* (*X* = *x*) is the probability of each dose in each individual and *p* is the allele frequency. Assuming HWE, we can find the expected dosage variance for a particular dosage, given the expectation of *X*

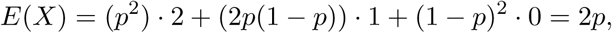

and the usual formula for the variance of a random variable to find the dosage variance:

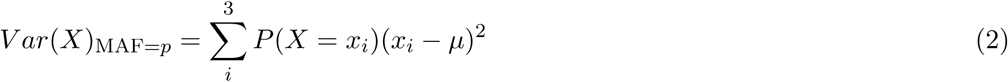

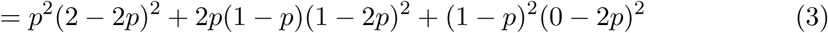

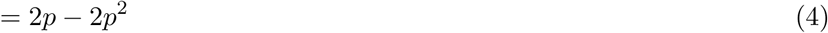

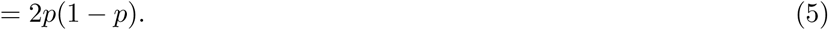

We assume that each SNP has an effect *β* and its estimated value *β̂* is estimated from the data using ordinary least squares. We assume a set of phenotypic associations, *Y* = *y_d_, y_r_,…, y_n_*, that can be modeled with a linear model of the SNP effect as follows:

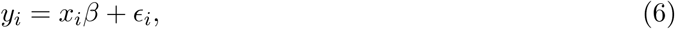

where *ɛ* is a noise term ∼ *N* (0*, σ*^2^), noting that *X* is distributed according to HWE as shown above for any particular allele frequency *p*. Using ordinary least squares, *β̂* would be estimated as follows:

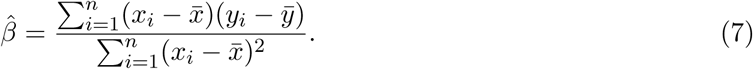

The variance of *β̂* is defined with an estimator 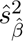 as follows:

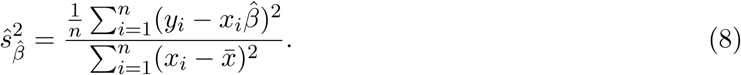

We want to know how *E*[*β̂*] and *E*[*ŝ_β̂_*] change with respect to *n* and *p*. First, let us solve *E*(*β̂*):

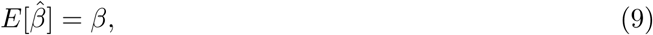

which is a well-known result. Next, consider the expectation of *ŝ_β̂_* with respect to *n* and *p*:

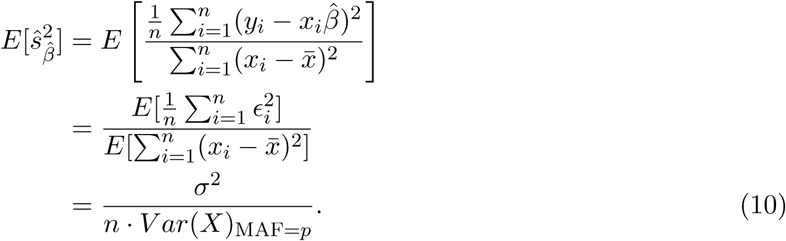

Thus, the expected slope estimate (*β̂*) is *β*, and does not depend on *p* or *n*. If we observe an arbitrary *β̂_d_* in one experiment, then in a repeated experiment the most likely value for *β̂_r_* is *β̂_d_* regardless of *p*. However, we can see that 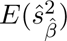 depends both on *n* and *p*. Therefore, if we observe QTL *β̂_d_* with MAF *p_d_* and number of observations *n_d_*, then if the same experiment is replicated at MAF *p_r_*, we expect:

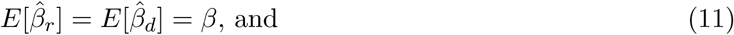

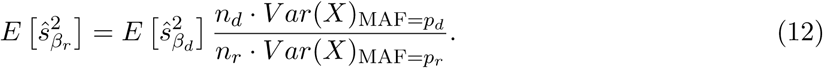

### Simulation

We tested this hypothesis in simulations, in which we repeatedly simulated *y* according to Eqn. (6), with *β* = 1*, σ* = 1, and *n* = 100 for values of 0 *< p <* 1, assuming the genotype is in HWE.

We then estimated the *β̂* and *s_β̂_* from the simulated data. We plot *β̂* and *s_β̂_* (shown in blue) in terms of *p*, as well as *E*(*β̂*) and *E*(*s_β̂_*) from Eqns. (9,10) (shown in red). From Figure SN1 we can see the observational data matches our theoretical expectations. Moreover, the red lines show us how the expected *β̂* and *s_β̂_* should change as the MAF changes between otherwise identical experiments.

**Figure SN1.**
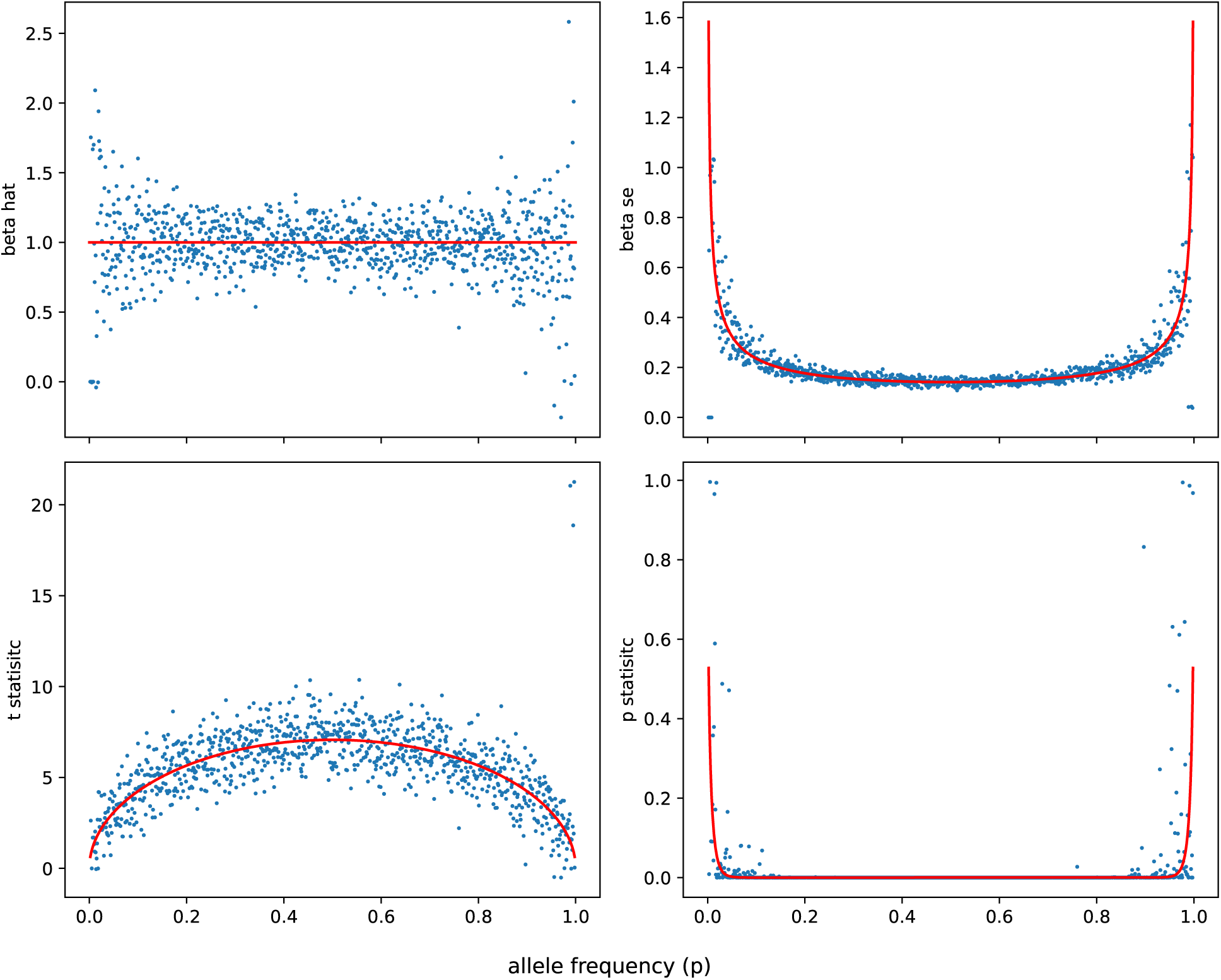
Sensitivity of effect estimation to allele frequency in GWAS experiments. Each panel plots a statistic from ordinary-least-squares estimation of a simulated QTL against the variant’s allele frequency (x-axis, 0 *< p <* 1). Data were generated under Eqn. (6) with *β* = 1, *σ* = 1, and *n* = 100 at each allele frequency, assuming the genotype is in HWE. Blue points show values estimated from the simulated data; red lines show the theoretical expectations from Eqns. (9,10). **beta hat** (*β̂*) is the estimated effect size, **beta se** (*s_β̂_*) its standard error, **t statistic** is *β̂/s_β̂_*, and **p statistic** is the corresponding *p*-value. The estimated effect size stays centred on the true value *β* = 1 across the whole frequency range, whereas its standard error rises steeply as variants become rarer (*p* → 0 or 1), because the genotype variance 2*p*(1 − *p*) shrinks; consequently the *t*-statistic falls and the *p*-value grows at low MAF.

### Simulation on 1000 Genomes Dataset

We also simulated eQTL data using the 1000 genomes dataset (1KG) to test this method [48–50]. In each of the 1KG super populations (EUR, AMR, SAS, AFR) we extracted SNPs within 1 mega-base from the transcription start site of protein-coding genes in the human genome. Then for each extracted gene we simulated a quantitative trait with one causal variant with the following equation:

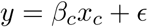

where the effect size *β_c_* was set to one and *ɛ* ∼ *N* (0, 3). When synthetically producing these summary statistics we down-sampled populations to the following numbers to capture a range of population sizes and ancestries: EUR n=500, AFR n=500, AMR n=250 and SAS n=100. We compared the European ancestry summary statistics to each of the other three populations, we are able to predict 73% of portable lead eQTLs (Figure SN2).

**Figure SN2.**
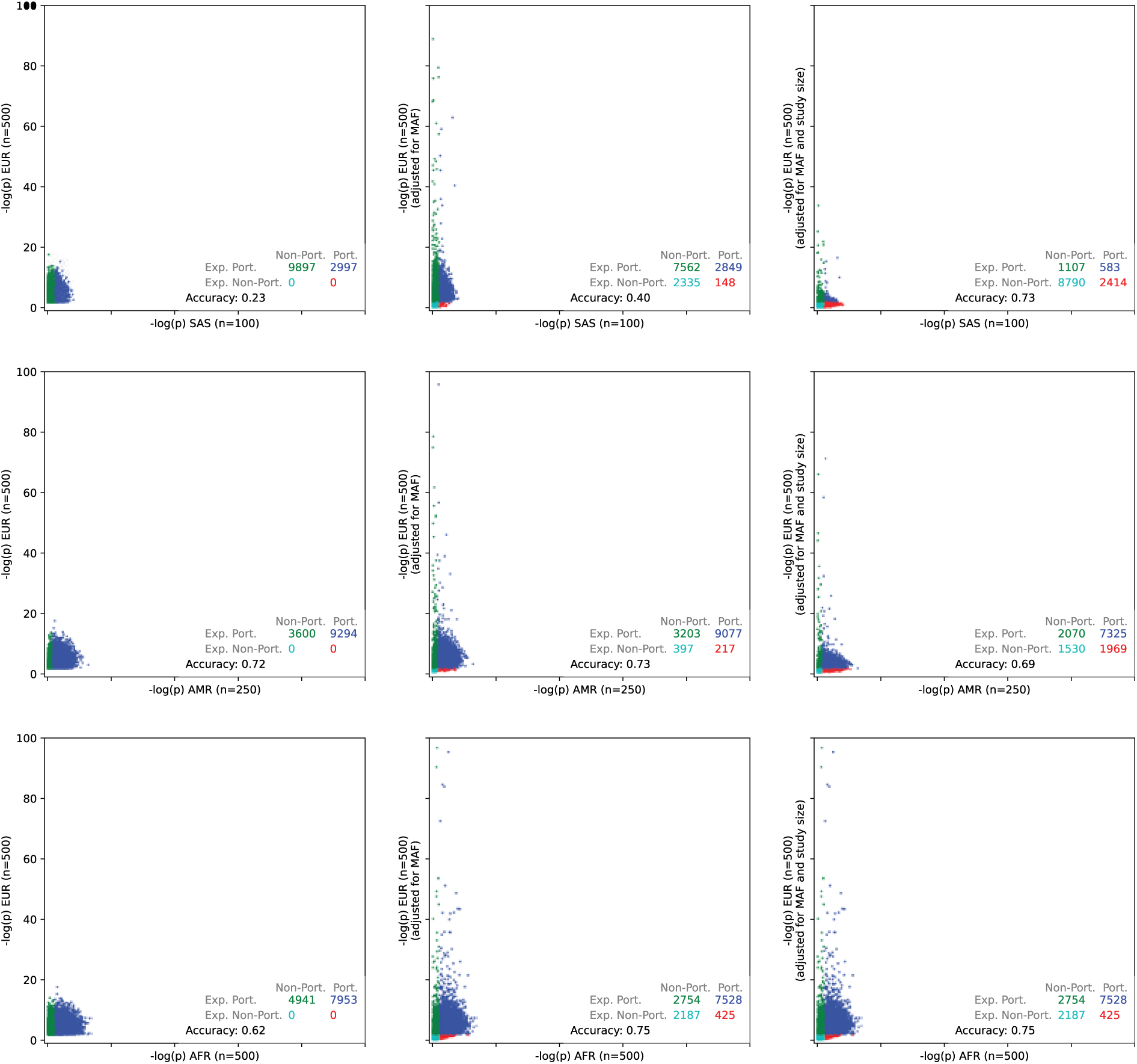
Accounting for sample-size and allele-frequency differences recovers portable eQTLs across 1000 Genomes populations. Quantitative traits were simulated from real 1000 Genomes genotypes in four super-populations (EUR, AFR, AMR, SAS), using variants within 1 Mb of the transcription start site of each protein-coding gene. For every gene a single causal variant was assigned effect *β_c_* = 1 with residual noise *ɛ* ∼ *N* (0,3), and populations were down-sampled to span a range of sizes and ancestries (EUR *n* = 500, AFR *n* = 500, AMR *n* = 250, SAS *n* = 100). Taking EUR as the discovery sample and each remaining population as the replication sample, the framework predicts the replication effect size and standard error from the discovery estimates together with the two populations’ sample sizes and allele frequencies (Eqns. (11,12)).

## Supplementary Material

Supplementary materials include 13 supplementary figures and 8 supplementary tables.

## Supplementary figures

**Figure S1 *coloc-SuSiE* LD diagnostic plots** Each figure is a diagnostic plot showing the consistency between summary statistics and the reference LD matrix, based on the residual sum of squares model under the null with a regularised LD matrix. Each subplot is shown for 10 arbitrary eGenes (columns) across all populations in the study (rows). The x-axis shows the expected z-statistic inferred using the LD matrix, and the y-axis shows the observed z-statistic.

**Figure S2 The number of eSNP and eGene discoveries is log-linearly correlated with the number of observations in each study.** y axis shows the log sample size for each cohort, x axis shows the number of eSNP/eGene discoveries

**Figure S3 Portability of lead eQTLs (with FDR *<* 0.05) across different SNP-level statistical significance and effect ratio thresholds.** (A) Uncertainty unaware effect ratio, portable if the ratio of the marginal effect sizes is within the specified range. (B) Effect ratio adjusted by MASHR. (C) uncertainty-aware effect ratio, portable if the 95% CI of the effect ratio overlaps the specified range. (D) Portable if a lead eSNP is also significant in the replication cohort at a given FDR threshold. **Study Set 1** Portability between African-American, European-American and Hispanic cohorts in CD14^+^ monocytes. **Study Set 2** Portability between Puerto Rican, African-American and Mexican-American cohorts in whole blood. **Study Set 3** Portability between Indonesian and three sets of European and European-American cohorts in whole blood eQTLs. For all studies, we use an *FDR <* 0.05 threshold to identify eQTLs in the discovery cohort, then vary the statistical significance threshold applied to the replication cohort as shown in the figure.

**Figure S4 Portability of all eQTLs (with FDR *<* 0.05) across different SNP-level statis-tical significance and effect ratio thresholds.** (A) Uncertainty unaware effect ratio, portable if the ratio of the marginal effect sizes is within the specified range. (B) Effect ratio adjusted by MASHR. (C) uncertainty-aware effect ratio, portable if the 95% CI of the effect ratio overlaps the specified range. (D) Portable if a eSNP is also significant in the replication cohort at a given FDR threshold. **Study Set 1** Portability between African-American, European-American and Hispanic cohorts in CD14^+^ monocytes. **Study Set 2** Portability between Puerto Rican, African-American and Mexican-American cohorts in whole blood. **Study Set 3** Portability between Indonesian and three sets of European and European-American cohorts in whole blood eQTLs. For all studies, we use an *FDR <* 0.05 threshold to identify eQTLs in the discovery cohort, then vary the statistical significance threshold applied to the replication cohort as shown in the figure.

**Figure S5 Portability of eQTLs across different gene-level significance thresholds.** (A) Portable to the replication cohort if at least one SNP has an FDR-adjusted p-value less than the specified threshold. (B) Portable to the replication cohort if the two genes colocalize with a CCV above the specified threshold. **Study Set 1** Portability between African-American, European-American and Hispanic cohorts in CD14^+^ monocytes. **Study Set 2** Portability between Puerto Rican, African-American and Mexican-American cohorts in whole blood. **Study Set 3** Portability between Indonesian and three sets of European and European-American cohorts in whole blood eQTLs. For all studies, we use an *FDR <* 0.05 threshold to identify eQTLs in the discovery cohort, then vary the gene-wise statistical significance or CCV threshold applied to the replication cohort as shown in the figure.

**Figure S6 Sample size and eQTL portability** (A-F) The relationship between population size and portability The y-axis of subplot shows the mean number of eQTLs from a discovery cohort, that are portable to a second study cohort. The x-axis shows number of individuals in replication cohort. Each point represents every possible pair of populations from each Study Set. Colour and shape is defined by the genetic ancestry and study cohort of replication cohort for each of these pairs. **(A)** Proportion of eSNPs from the discovery cohort that have an effect ratio *>* 0.5 in replication cohort. **(B)** Proportion of eSNPs from the discovery cohort that have a *FDR >* 0.05 in replication cohort. **(C)** Proportion of eGenes portable as defined by having at least 1 eSNP in both the discovery cohort and replication cohort. **(D)** Proportion of lead eSNPs from the discovery cohort with an effect ratio *>* 0.5 in replication cohort. **(E)** Proportion of lead eSNPs from the discovery cohort portable that have a *FDR <* 0.05 in replication cohort. **(F)** Proportion of eGenes portable defined as colocalising with CCV *>* 0.5 in the ‘coloc‘ R package.

**Figure S7 Predictors of replication p-values among lead eSNPs of the discovery population** (A–C) Show Study Sets 1–3, respectively. Each cell in the heatmap represents the Spearman correlation coefficient between the nominal p-values in the replication population and SNP portability predictors (among lead eSNPs of the discovery population). “t-stat” refers to the SNP t-statistic; MAF refers to minor allele frequency; and gnomAD refers to SNP LD scores across four super populations: AFR, AMR, EAS, and NFE.

**Figure S8 Predictors of differences in log p-values between discovery and replication populations among lead eSNPs of the discovery population** (A–C) Show Study Sets 1–3, respectively. Each cell in the heatmap represents the Spearman correlation coefficient between the difference in log(P) values between the discovery and replication populations (among lead eSNPs of the discovery population) and SNP portability predictors. “t-stat” refers to the SNP t-statistic; “MAF” refers to SNP minor allele frequency; and “gnomAD” refers to SNP LD scores across four super populations: AFR, AMR, EAS, and NFE.

**Figure S9 Predictors of effect size ratios between discovery and replication populations among lead eSNPs of the discovery population** (A–C) Show Study Sets 1–3, respectively. Each cell in the heatmap represents the Spearman correlation coefficient between the effect size ratio of the replication population to the discovery population (among lead eSNPs of the discovery population) and SNP portability predictors. “t-stat” refers to the SNP t-statistic; “MAF” refers to SNP minor allele frequency; and “gnomAD”refers to SNP LD scores across four super populations: African (AFR), Admixed American (AMR), East Asian (EAS), and Non-Finnish European (NFE).

**Figure S10 Predictors of replication p-values among all eSNPs of the discovery population** (A–C) Show Study Sets 1–3, respectively. Each cell in the heatmap represents the Spearman correlation coefficient between the nominal p-values in the replication population and SNP portability predictors (among all eSNPs of the discovery population). “t-stat” refers to the SNP t-statistic; “MAF” refers to SNP minor allele frequency; and “gnomAD” refers to SNP LD scores across four super populations: AFR, AMR, EAS, and NFE. (D) The y-axis shows, for each discovery–replication population pair, the correlation between differences in SNP variance and differences in log(P) values among all eSNPs of the discovery population, with the x-axis showing the sample size of the replication population.

**Figure S11 Predictors of differences in log p-values between discovery and replication populations among all eSNPs of the discovery population** (A–C) Show Study Sets 1–3, respectively. Each cell in the heatmap represents the Spearman correlation coefficient between the difference in log(P) values between the discovery and replication populations (among all eSNPs of the discovery population) and SNP portability predictors. “t-stat” refers to the SNP t-statistic; “MAF” refers to SNP minor allele frequency; and “gnomAD” refers to SNP LD scores across four super populations: AFR, AMR, EAS, and NFE. (D) The y-axis shows, for each discovery–replication population pair, the correlation between differences in SNP variance and differences in log(P) values among all eSNPs of the discovery population, with the x-axis showing the sample size of the replication population.

**Figure S12 Predictors of effect size ratios between discovery and replication populations among all eSNPs of the discovery population** (A–C) Show Study Sets 1–3, respectively. Each cell in the heatmap represents the Spearman correlation coefficient between the effect size ratio of the replication population to the discovery population (among all eSNPs of the discovery population) and SNP portability predictors. “t-stat” refers to the SNP t-statistic; “MAF” refers to SNP minor allele frequency; and “gnomAD” refers to SNP LD scores across four super populations: AFR, AMR, EAS, and NFE. (D) The y-axis shows, for each discovery–replication population pair, the correlation between SNP variance and effect size ratio among all eSNPs of the discovery population, with the x-axis showing the sample size of the replication population.

**Figure S13 SNP portability explained by SNP features by the size each study** y-axis shows the *r*^2^ score between a portability metric from a discovery cohort and replication cohort (rows). x axis shows the sample size of each study. **(A-F)** shows correlations among lead eSNPs. **(G-L)** shows correlations among all significant eSNPs.

## Supplementary tables

**Table S1 Summary metadata, including exclusion reason where relevant, for all studies we considered including in our analysis.**

**Table S2 Portability between all cohorts at multiple statistical significance and effect size ratio cutoffs.** eQTLs are identified in the discovery cohort using a global FDR *<* 0.05 threshold, and then ported to varying replication cohorts as described in the text.

**Table S3 Jaccard similarity between eQTL measures across each Study Set.**

**Table S4 Jaccard similarity between eQTL measures comparing different effect ratio cuttoffs to the statistical significance portability metric of FDR** *<* **0.05.**

**Table S5 Adjusted portability results for lead eSNPs in all three Study Sets after correcting for MAF and sample size differences.** “None” indicates typical portability. “MAF adjustment” refers to adjusting the p-statistic in the discovery cohort to match the SNP variance of the replication cohort. “Size adjustment” refers to adjusting the p-value in the discovery cohort to match the sample size of the replication cohort. **TP**: eSNPs expected to be portable and were portable in the empirical data after adjusting the replication cohort. **FP**: eSNPs expected to be portable and were non portable in the empirical data after adjusting the replication cohort. **FN**: eSNPs expected to be non portable and were portable in the empirical data after adjusting the replication cohort. **TN**: eSNPs expected to be portable and were non portable in the empirical data after adjusting the replication cohort. **Accuracy**: proportion of correctly classified eSNPs.

**Table S6 eSNP portability between cohorts, comparing MASH and FDR derived significance tests for eSNPs.** Tables show cohorts pooled with MASH or fixed effect meta-analysis, and how well they are predicted by eSNPs that are significant (by FDR) in a held out cohort.

**Table S7 eGene portability between cohorts, comparing MASH and FDR derived significance tests for eGenes.** Tables show cohorts pooled with MASH or fixed effect meta-analysis, and how well they are predicted by eGenes that are significant (by FDR) in a held out cohort.

**Table S8 lead eSNP portability between cohorts, comparing MASH and FDR derived significance tests for lead eSNPs.** Tables show cohorts pooled with MASH or fixed effect meta-analysis, and how well they are predicted by lead eSNPs that are significant (by FDR) in a held out cohort.

